# Profiles of secoiridoids and alkaloids in tissue of susceptible and resistant green ash progeny reveal patterns of induced responses to emerald ash borer in *Fraxinus pennsylvanica*

**DOI:** 10.1101/2022.05.18.492370

**Authors:** Robert K. Stanley, David W. Carey, Mary E. Mason, Therese M. Poland, Jennifer L. Koch, A. Daniel Jones, Jeanne Romero-Severson

## Abstract

The emerald ash borer (*Agrilus planipennis*, EAB) invasion in North America threatens most North American *Fraxinus* species, including green ash (*F. pennsylvanica*), the mostly widely distributed species (1, 2). A small number of green ash (“lingering ash”, 0.1-1%) survive years of heavy EAB attack (3) and kill more EAB larvae when challenged in greenhouse studies than susceptible controls (4). We combined untargeted metabolomics with intensive phenotyping of segregating F_1_ progeny from susceptible or lingering ash parents to detect chemotypes associated with defensive responses to EAB. We examined three contrasting groups: low larval kill (0-25% of larvae killed), high larval kill (55-95% of larvae killed) and uninfested. Contrasting the chemotypes of these groups revealed evidence of an induced response to EAB. Infested trees deployed significantly higher levels of select secoiridoids than uninfested trees. Within the infested group, the low larval kill (LLK) individuals deployed significantly higher levels of select secoiridoids than the high larval kill (HLK) individuals. The HLK individuals deployed significantly higher concentrations of three metabolites annotated as aromatic alkaloids compared to the LLK and uninfested individuals. We propose a two-part model for the North American *Fraxinus* response to EAB wherein every individual has the capacity to detect and respond to EAB, but only certain trees mount an effective defense, killing enough EAB larvae to prevent or minimize lethal damage to the vascular system. Integration of intensive phenotyping of structured populations with metabolomics reveals the multi-faceted nature of the defenses deployed in naïve host populations against invasive species.

**Significance:** Long-lived forest trees employ evolutionarily conserved templates to synthesize an array of defensive metabolites. The regulation of these metabolites, honed against native pests and pathogens, may be ineffective against novel species, as illustrated by the high mortality (>99%) in green ash infested by the invasive emerald ash borer (EAB). However, high standing genetic variation may produce a few individuals capable of an effective defense, as seen in the rare surviving green ash. In an investigation of this plant-insect interaction, we annotated metabolites associated with generalized but ineffective responses to EAB, and others associated with successful defensive responses. Untargeted metabolomics combined with intensive phenotyping of structured populations provides a framework for understanding resistance to invasive species in naïve host populations.

## INTRODUCTION

Invasive pests and pathogens, now widely dispersed through globalization, threaten nearly two thirds of North American forests (1). Exacerbated by climate change, these increasingly severe infections and infestations destabilize forested ecosystems and inflict billions of dollars in direct costs to individuals and local communities (5-14). Two of these pathogens and pests, chestnut blight (*Cryphonectria parasitica*) and emerald ash borer (*Agrilus planipennis*, EAB) have had a profound impact on public awareness: chestnut blight because this disease caused the ecological extinction of the iconic American chestnut (*Castanea dentata*) and EAB because of the widespread, rapid and continuing loss of ash trees from streets, parks, and forests (14). The severe impact of the loss of chestnut, pales in comparison to the economic and ecological damage already inflicted by EAB, the most destructive and economically devastating invasive insect pest of forest trees in North American history (15).

EAB, a beetle native to Asia, was discovered in Michigan, United States and in Ontario, Canada in 2002 (16). EAB attacks ash (*Fraxinus*) species; larvae hatch from eggs laid in bark furrows and burrow into the living tissue directly beneath the bark (2). The larvae feed on the vascular cambium, cork cambium, phloem, and xylem inflicting severe vascular damage that ultimately kills the tree. Larvae feed during the summer and the fall and may take one or two years to complete development. The larvae pass through four developmental instars and then chew a pupal chamber in the outer sapwood or inner bark in which they overwinter as mature larvae folded over in a J-shape. During the spring, they enter the pupal stage and transform into adults that emerge in late spring and early summer through characteristic D-shaped exit holes (17, 18).

EAB infestation has resulted in the rapid loss of hundreds of millions of green ash, not only in forests and rural areas but also in cities, where green ash was once one of the most widely planted street and park trees in the United States (19, 20). Green ash, the most widely distributed *Fraxinus* species in North America, is a dioecious, diploid, and deciduous tree species native to the eastern and central United States and eastern Canada (16, 18, 20). Mortality in green ash from EAB infestation can approach 100% within six years of the local detection of EAB (20). EAB invasion threatens not only green ash, but survival of the majority of native North American *Fraxinus* species including white (*F. americana*), pumpkin (*F. profunda*), Carolina (*F. caroliniana*) and black ash (*F. nigra*) (20, 21). *Fraxinus* are ecologically important in a wide range of forested ecosystems and are also extensively utilized for soil conservation, rural water management, riparian zone stabilization, flood control, and urban green spaces in North America (19).

Long term forest plot monitoring initiated in 2005, two years (3, 20, 22) after the initial detection of EAB in North America, revealed a small number of green ash (0.1-1%) that survive for years after all other surrounding green ash have died (3). These “lingering ash” (L) have been and continue to be propagated as potential sources of genetic resistance for a breeding program (23, 24). Eleven years of replicated egg bioassay tests, conducted by placing controlled densities of EAB eggs on test trees and monitoring larval development and survival, revealed reproducible larval kill capabilities with phenotypic distributions among trees that suggest quantitative inheritance (24, 25).

Durable genetic resistance in the host, the most effective control measure for any pest or pathogen (26, 27), was not initially considered a strategic goal for saving North American *Fraxinus* species from EAB. The assumption was that a species cannot have any resistance to a pest with which it has not coevolved (28, 29). However, many studies have shown that in many cases native species do marshal heritable defensive responses to non-native invaders (30, 31). Successful breeding programs have produced American *beech* (*Fagus grandifolia*) resistant to beech bark disease (*Neonectria spp* transmitted by *Cryptococcus fagisuga*) (32), eastern white pine (*Pinus strobus*) resistant to white pine blister rust (*Cronartium ribicola*) (33) and Port Orford cedar (*Chamaecyparis lawsoniana*) (22) resistant to the root rot pathogen *Phytophthora lateralis*. The success of these and other programs demonstrates that heritable resistance exists in wild populations and can be used to develop resistant populations for species restoration through breeding (30). Once a genetic component is confirmed, a detailed study of the mechanisms of the response can contribute to a body of knowledge on the omics of heritable defensive responses.

Previous investigations on the role of secondary metabolites as defenses against EAB have focused on comparing small numbers of cultivars from susceptible *Fraxinus spp*. to the naturally resistant *F. mandshurica* cultivar *‘mancana’*(34). Application of methyl jasmonate in infested susceptible *F. americana* individuals induced production of verbascoside and suppressed EAB larval development (35).These studies collectively proposed a positive association between lignan glycosides and host resistance, particularly pinoresinol and verbascoside, as well as suggesting a role for secoiridoid glycosides (36).

Secoiridoids are also implicated in the response of European Ash (*F. excelsior*) to ash dieback disease (ADB) caused by *Hymenoscyphus fraxineus*. Ash dieback ultimately infects the woody stem tissue, killing the tree (37, 38). High concentrations of specific secoiridoids were identified with tolerant genotypes in one study, and with susceptible genotypes in another. Both groups of investigators proposed that the different levels of secoiridoids are the result of differential transcriptional regulation (39, 40). Investigations of ash dieback phenotypes, in these and other studies suggest that susceptibility to ADB in *F. excelsior*, is a quantitative trait (41).

Other recent investigations of resistance to wood-boring insects have shown that some trees utilize secondary metabolite-based constitutive and induced defensive responses against specific insect pests (42-44). The concentration and profiles of certain plant secondary metabolites strongly predict resistance in maritime pine *(Pinus pinaster*) to the pine weevil (*Hylobius abietis*), after accounting for genetic relatedness among the host trees (42). Other investigations have shown that the response consists of altered rates of synthesis for existing metabolites, rather than the synthesis of unique compounds (42, 45).

Here we combine untargeted metabolomics and intensive phenotyping on structured populations using an experimental design that accounts for the confounding effect of genetics and environment to detect chemotypes associated with defensive responses to EAB. We hypothesized that the full sibling progeny of Susceptible x Susceptible (SxS), and Lingering x Lingering (LxL) parents would produce a wide range of larval kill phenotypes and that the family means of the progeny from LxL parents would be significantly higher than the family means of the progeny of SxS parents. If both these hypotheses are correct, and the defense is associated with secondary metabolites, we expect a contrast in chemotypes between the high larval kill (HLK, tree defenses killed 55-95% of larvae) and low larval kill (LLK, tree defenses killed 0-25% of larvae) phenotypes. If infestation induces a response, we expect that the chemotypes of infested individuals will be distinct from uninfested individuals within families. Our data showed that some secondary metabolites including select secoiridoids occurred at higher concentrations in infested individuals regardless of larval kill phenotype, while a smaller number of compounds, annotated as aromatic alkaloids were found in higher concentrations in high percent larval kill individuals. Our work will spur future investigations for the molecular basis of durable genetic resistance to EAB in green ash and provide a framework for discovering resistance to invasive species in naïve host populations.

## RESULTS

### Analysis of EAB-resistance in full-sibling families of reveals that resistance to EAB in green ash is a multigenic quantitative trait

Seedlings (2-3 years old) from two green ash F_1_ families produced through crosses between lingering parents (LxL) and one family produced by a cross between susceptible parents (SxS) were infested with EAB to confirm the genetic basis of the larval kill phenotype (Figure 1a).. One-way ANOVA and Tukey-Kramer multiple comparison tests revealed that the mean percent larval kill of (LxL) families Pe-Y and Pe-Z were significantly different from the mean percent larval kill of the (SxS) family Pe-C (p < 0.01), but there was no significant difference among the L x L families’ means (Figure. 1b). The shape and range of the larval kill distributions strongly suggests that the phenotype is a quantitative trait and provides support for the hypothesis of complex inheritance (Figure. 1b).

**Figure 1:**
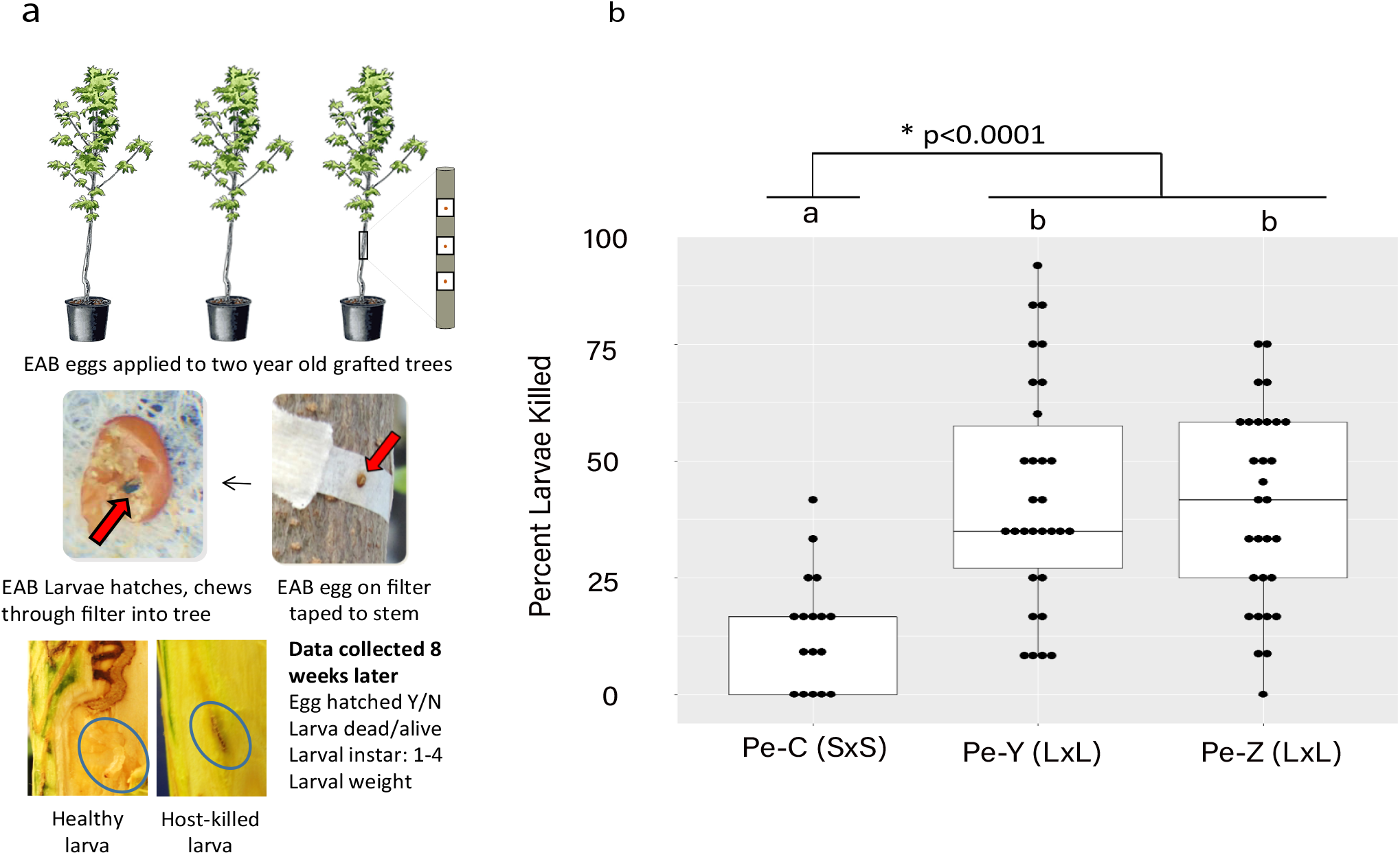
Egg Bioassay Protocol & Phenotypic Distributions. (a) EAB greenhouse bioassay protocol. Two year old green ash individuals are artifcially dissected at eight weeks to ascertain larval fate. (b) Percentage of EAB larvae kill by infested individuals in families Pe-C, Pe-Y and Pe-Z when sampled 8 weeks post infestation. Family C contained 17 F_1_ infested progeny from two susceptible parents. Families Pe-Y and Pe-Z both contained 30 F_1_ full sib progeny of two lingering ash parents. Family means of Pe-Y and Pe-Z were significantly different from the family mean of susceptible family C (p<0.0001)

Each family produced a range of larval kill phenotypes, (Pe-C: 0-44%, Pe-Y: 8-95%, Pe-Z: 0-75%). Based on the distribution of phenotypes across families (Figure. 1b), we classified individual trees with larval kill values of 25% or lower as LLK and those with larval kill values greater than 55% as HLK. The value of 55% is higher than the highest larval kill value for the collection of lingering ash parents described in a previous report, and the value of 25% is higher than the parents of family Pe-C, and most of the progeny (88%) in the susceptible family Pe-C (Figure 1) (24). As a comparison, the resistant Asian ash *F. mandshurica* typically kills 80-90% of EAB larvae when tested with the egg bioassay (24). The lingering families in this study included some progeny that performed similarly to resistant Manchurian ash individuals.

### Generation of untargeted metabolomic profiles

We produced untargeted metabolomic profiles from acetonitrile:isopropanol:water extractions using ultra-high performance liquid chromatography/ high resolution mass spectrometry (UHPLC/MS). The levels of metabolites were normalized to a constant internal standard and replicated, with a constant mass of tissue extracted. An analysis of the relative standard deviation (RSD) of pooled controls had a median of 29.8% for all features considered in downstream analyses (Figure S1)

### Metabolite based OPLS-DA models correctly identify progeny classes

We conducted pairwise comparisons of the metabolite profiles of HLK, LLK, and uninfested (UNI) individuals within families to determine if metabolites were associated with infestation status or the larval kill phenotype. We assessed 194 metabolite features (Figure 2) with pairwise one-way analysis of variance (ANOVA) tests for the following contrasts: Family C: UNI vs LLK; Family Y: UNI vs LLK, UNI vs HLK, LLK vs HLK; Family Z: UNI vs LLK, UNI vs HLK, LLK vs HLK. Between 9 and 49 features were significant (p < 0.05) in each comparison (Figure 2, Table S2).

**Figure 2:**
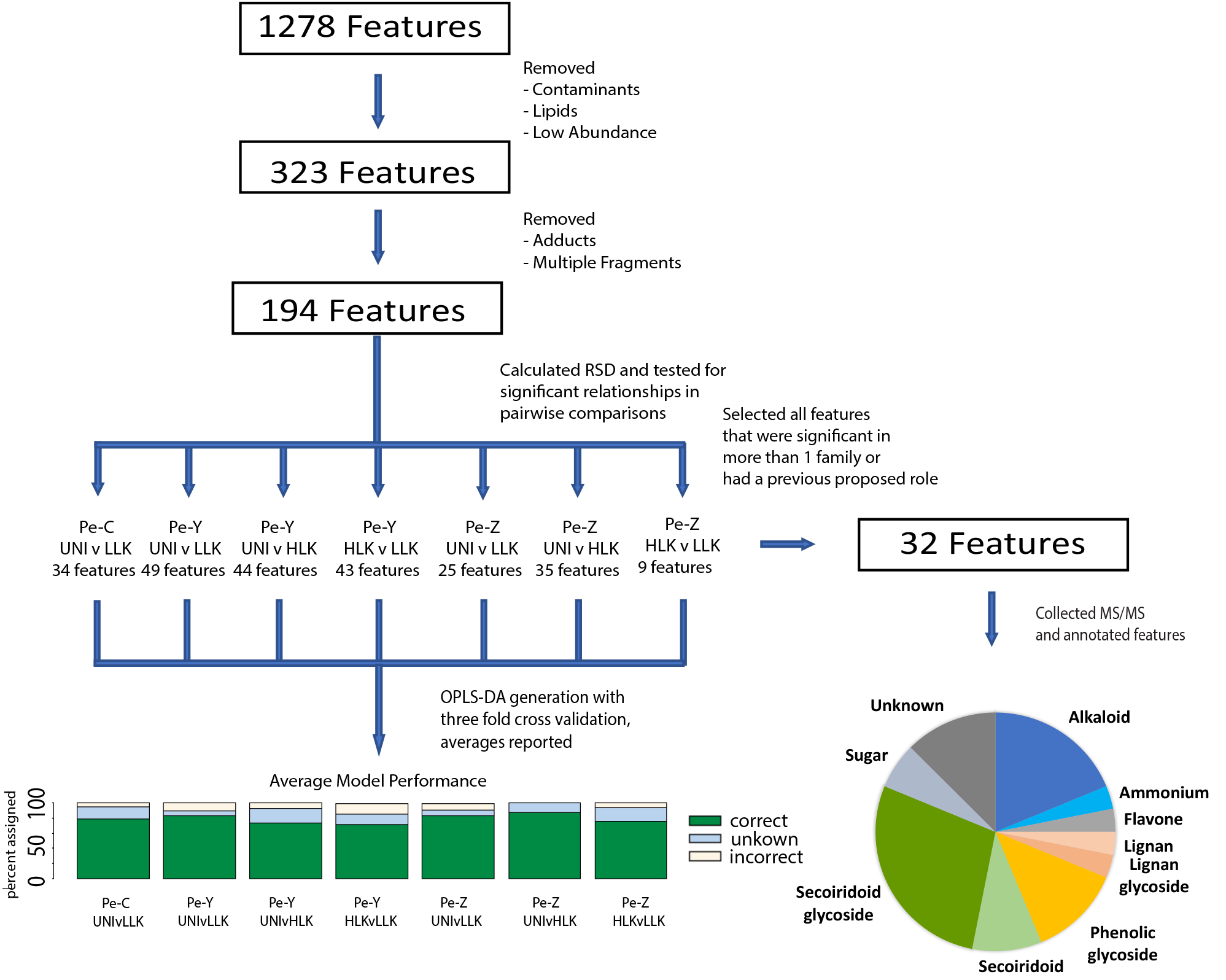
Data Processing Schematic. Flowthrough of the untargeted metabolomics workflow beginning with data generation, and highlighting the number of features at each stage of the analysis.

We then took these features and used orthogonal partial least squares-discriminant analyses (OPLS-DAs) to examine their ability to accurately identify the correct progeny classification (Figure 2). To prevent overfitting of our model we performed a three-fold cross validation on our data and report the average prediction accuracies as the performance of our model. Overall, our model performed quite well, with over 70 % of individuals correctly assigned to progeny larval-kill phenotype class across all models, with the majority of other individuals being unclassified, not incorrectly assigned (Figure 3a, Table S2). Our confidence in these models was supported by principal component analyses (PCAs) yielding similar separations(Figures 3b, 3c), suggesting that OPLS-DAs are producing statistically meaningful group separations (46). This workflow can serve as a template for assessing the relationship of chemotypes and complex phenotypes in a non-model system.

**Figure 3:**
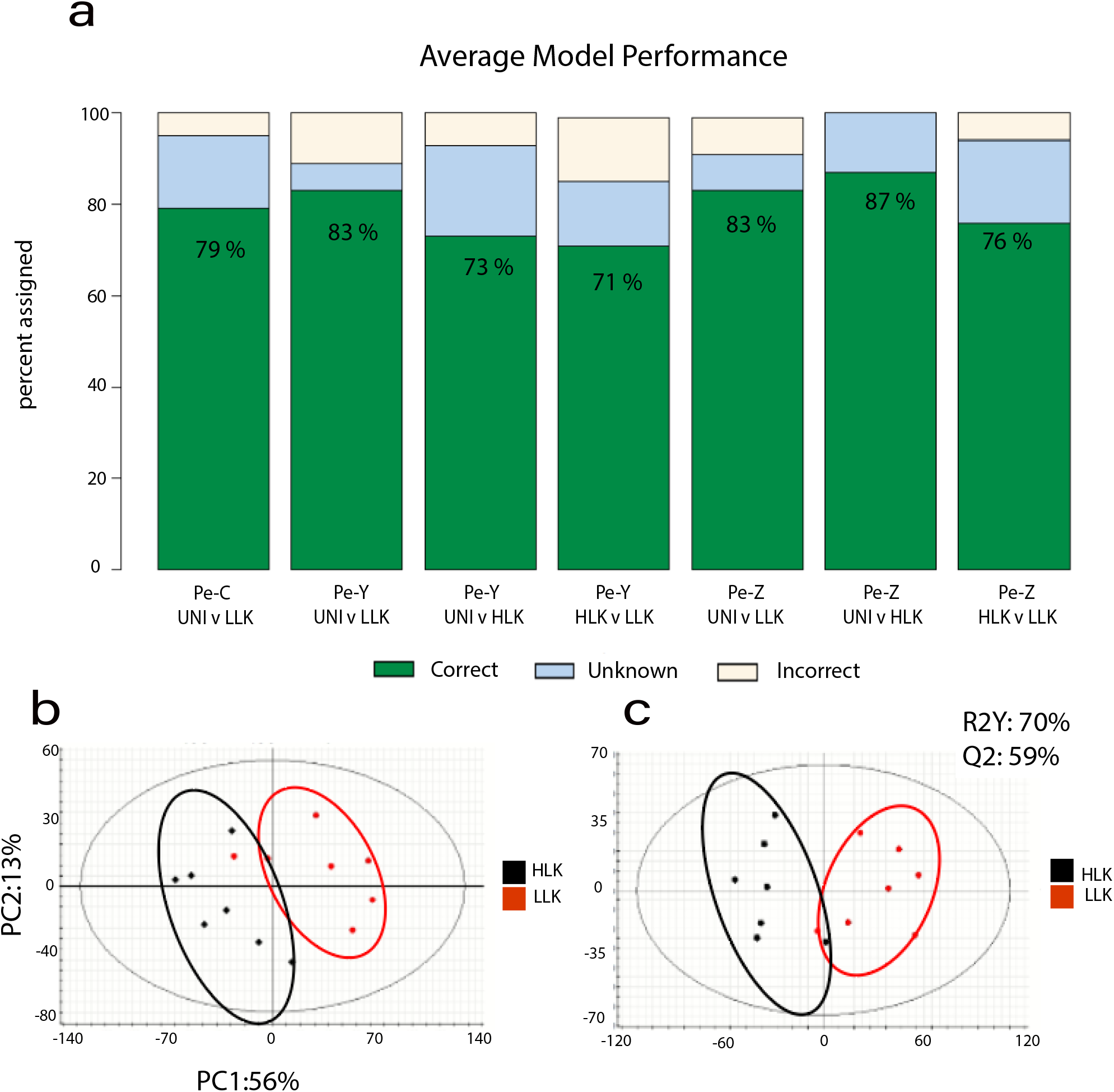
Classification summary. (a) Orthogonal partial least squares projection to latent square discriminate analysis (OPLS-DA) model performance, averaged across triplicate prediction models. The graph indicates that percentage that each model classified correctly, incorrection, or was unable to classify. (b) Principal component analysis plot comparing high and low larval kill (LK) in family Pe-Y, utilizing 43 features. (c) OPLS-DA model utilizing all test samples in a comparison of high vs low larval kill using 43 features.

### Chemotypes across families distinguish a general defense response from a successful defense response

We focused our attention on the 32 features that had a significant *p*-value in more than one family’s comparisons or were suggested as important for EAB defense in previous investigations. This latter category included verbascoside (35) and salidroside (47). We generated electrospray ionization tandem mass spectra (ESI-MS/MS) for the features detected in our analysis and annotated them based on comparisons with MS/MS databases including the Massbank of North America, along with published literature and purchased standards. The annotation confidence is labeled according to the recommendations of the Metabolomics Standards Initiative (MSI) (48). Our annotations revealed compounds from a wide variety of chemical families, including the first record of specific alkaloids present in green ash tissue (Table 1, Figure 4, Figure 5).

**Figure 4:**
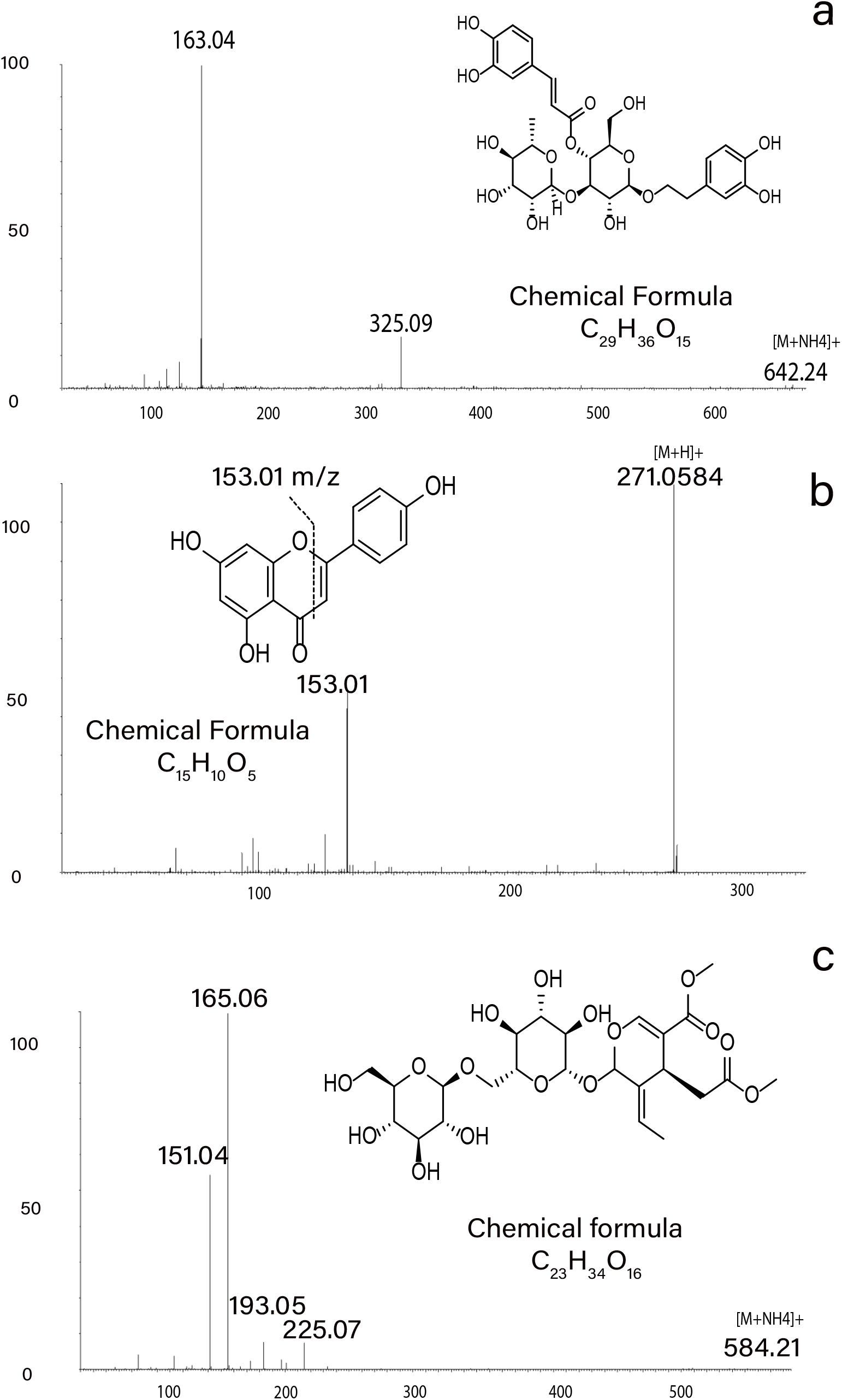
Metabolite Annotations. MS/MS spectra in positive ion mode support annotations of metabolite structures: (a) product ions of m/z 642.24 ([M+NH_4_]^+^) for verbascoside, (product ions of m/z 271.06 ([M+H]^+^) for apigenin, (c) product ions of m/z 584.21 ([M+NH_4_]^+^) for Excelside A.

**Figure 5:**
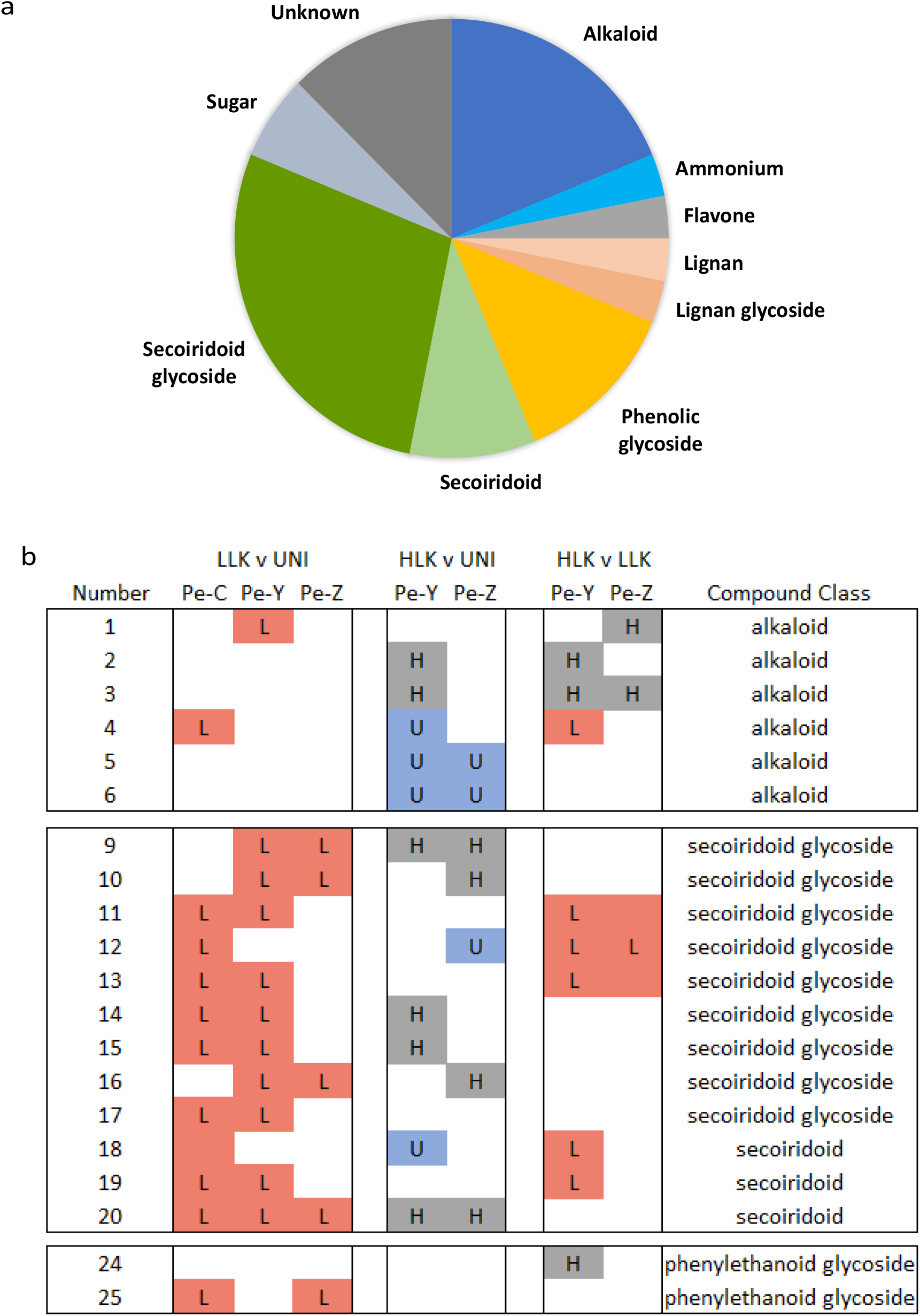
Chemical Families of Annotated Compounds. (a) proportions of the chemical families in the annotated metabolites. (b) Pairwise comparisons for specific compounds. ‘Number’ is metabolite number (table 1). LLK v UNI, HLK v UNI, HLK v LLK indicates pairwise comparisons between larval kill phenotypes or uninfested individuals. Pe-C, Pe-Y, Pe-Z refer to full sibling families (figure 1). Box with letter indicates the phenotypic category that had significantly higher concentration of the indicated metabolite (p < 0.05, L in red LLK, H in gray HLK, U in blue UNI). Annotated metabolites 1-6 are alkaloids, 9-17 are secoiridoid glycosides, 18-20 are secoiridoids, 24 is salidroside and 25 is verbascoside.

We annotated 12 secoiridoids with similar structures to secoiridoids previously hypothesized to be indicative of a resistance mechanism (36). We found that these secoiridoids were elevated in both low and high larval kill phenotypes compared to uninfested controls (Figure S2). However, five secoiridoids had significantly higher concentrations in low larval kill phenotypes compared to high larval kill phenotypes with no significant difference in concentration in the other seven secoiridoids (Figure 5b-d).

One secoiridoid (*m/z 569*.*23*) was elevated in all infested comparisons (LLK v UNI, HLK v UNI) across all families, and may have some value as an indicator of infestation across many ash genotypes. Another two secoiridoids, nueznehide (*m/z 704*.*2781*) and GL5 (*m/z 928*.*3429*), found in higher concentrations in trees that are highly susceptible to ash dieback in previous investigations (39, 49), were significantly higher in low larval kill individuals compared to uninfested individuals across families. Additionally, we found that concentrations of verbascoside, a phenylethanoid glycoside also proposed as a component of the resistance response (36), were highest in low larval kill individuals. Overall, our data suggests that these specific secoiridoids and verbascoside may be indicative of a general wound response, but do not appear to be responsible for the high larval kill phenotype. The only compounds that were higher in high larval kill individuals compared to low larval kill individuals were three compounds annotated as aromatic alkaloids and the phenylethanoid glycoside salidroside (Table 1, Figure 5, Figure S3). These alkaloids are the first reported in green ash and suggest a novel role of alkaloid in defense against herbivory in forest trees.

## DISCUSSION

We investigated the ability of select green ash individuals to respond to EAB using structured populations, a reproducible phenotyping method, and an untargeted metabolomics approach. We found that all green ash seedlings analyzed displayed metabolic changes in response to infestation, but in most individuals, this response was ineffective to kill many EAB larvae. OPLS-DA and multivariate analyses showed that high and low performing individuals had chemotypes distinct from each other and from uninfested individuals. These chemotypes are distinguishable based on the relative concentrations of select metabolites (Table 2, Figure 5, Table S1), not their presence or absence, suggesting genetic regulation of multiple synthesis pathways may be responsible for the high larval kill phenotype. We provide an initial annotation of metabolites for further study, including secoiridoids that may prove to be reliable indicators of infestation across all genetic backgrounds, and three aromatic alkaloids that may be part of an effective defensive response.

Defensive responses based on multigenic mechanisms confer durable genetic resistance, the most effective control measure for any pest or pathogen. In our study, full sibling F_1_ progeny of lingering ash parents performed better on average than the F_1_ progeny of susceptible parents and produced progeny with phenotypes ranging from 0 % larvae killed to 95% larvae killed. This is the result expected when a phenotype is the result of complex genetic mechanisms involving multiple loci (50). A multigenic mechanism for the lingering ash phenotype is also consistent with two recent candidate gene studies, one on the pan-genome of EAB resistant *Fraxinus* species, and the other which utilized the 2021 release of the green ash genome (37, 51). Additional studies will be necessary to fully elucidate the genetic architecture of these defensive responses. Although we did not examine other components of the lingering ash phenotype, including adult feeding preferences or attractiveness of egg-laying sites to female EAB, our controlled greenhouse experiments did allow us to examine the defensive mechanisms deployed in the woody tissues, where the primary host insect interaction occurs.

The chemotypes of high larval kill, low larval kill, and uninfested individuals from the same parents could be distinguished based on relative concentrations of groups of metabolites. Comparisons of the same contrast across multiple families reveals that secoiridoids are associated with a generalized infestation response that does not predict effective defensive responses. This association is consistent with previous studies that suggested that infested trees, or trees artificially stressed with methyl jasmonate produced higher amounts of these metabolites (52). High concentrations of specific secoiridoids in *F. excelsior* are proposed to be indicative of tolerance (40) or susceptibility to ash dieback (37), and were predicted to provide a future robust reservoir of anti-feeding deterrents to EAB (49). Our data suggests that these specific secoiridoids function best as indicators of a generalized stress response and not necessarily of resistance to EAB. The study design allowed us to disentangle the generalized stress response from an effective defense response as indicated by percent larval kill. Our results suggest that part of the effective defensive response may consist of four metabolites, annotated as three aromatic alkaloids and salidroside, that were significantly elevated in high larval kill individuals compared to low larval kill or uninfested individuals. Overall, our study has distinguished, for the first time, between an effective defensive response and a generalized defensive response to EAB.

Based on our results, we propose a two-part model for the North American *Fraxinus* response to EAB wherein every individual has the biochemical capacity to synthesize chemical defenses as a response to EAB, but only certain trees deploy an effective induced defense response that kills enough EAB larvae to prevent or minimize lethal damage to the vascular system. This model is consistent with forest observations and controlled studies that show most individuals in North American ash species can kill a few larvae, but cannot withstand a heavy infestation (24, 25). The high concentrations of secoiridoid glycosides in infested individuals, especially those with the most live larvae, suggests that even susceptible ash trees detect that they have been wounded by EAB larvae, and attempt to respond but are unable to do so in a manner that results in effectively killing the larvae. A previous study demonstrated that application of methyl jasmonate induced a defensive response and suppressed EAB larval development or killed larvae in susceptible *Fraxinus* individuals (35), supporting the hypothesis that even susceptible trees have the necessary synthetic machinery, but lack the ability to conduct a tailored reconfiguration of their metabolism, as outlined by Schuman and Baldwin (53), to kill the EAB larvae.

This study provides a list of metabolites that could be targeted in future work focusing on the response of green ash to EAB. Key questions for future experiments include determining if the compounds identified extend to additional lingering ash families and gaining a better understanding of the timing and spatial distribution of effective defense responses. Additional phenotypic, genomic, proteomic, transcriptomic, and metabolomic analyses will benefit from the recent release of the green ash genome (51). This future work on the interaction of green ash and EAB will contribute to our understanding of how forest trees recognize and defend themselves against stem-boring insects.

In summary, our data supports the hypothesis that the high larval kill phenotype is a multi-genic and heritable trait. We have also shown that green ash responds to EAB infestation with increased concentrations of secoiridoids, regardless of the larval kill phenotype. While infestation with EAB induces a response in all green ash tested, the induced response is ineffective in most cases. In the individuals that mount a successful response, we found higher concentrations of three aromatic alkaloids and salidroside, a result that merits further investigation. Similar metabolites were seen across all phenotypes, but the concentrations varied, suggesting that the high larval kill phenotype is based on complex regulatory mechanisms. Elucidation of the genetic mechanisms driving defensive responses to EAB in green ash will be an essential part of a multidisciplinary effort for saving North American *Fraxinus* species and guide future investigations of resistance in native species to invasive threats.

## Materials & Methods

### Study System and Phenotyping

Green ash were selected in the forest based on two criteria: 1) a healthy canopy at least two years after the mortality rate of the stand exceeded 95 percent, and 2) a minimum diameter at breast height (DBH, 1.37 m from the ground) of 26 cm, indicating they were over the minimum size preferred by EAB when the infestation was at peak levels (24). These ‘lingering ash’ trees show evidence of less severe emerald ash borer infestation compared to susceptible phenotypes in the forest, often accompanied by evidence of vigorous wound healing, and maintain a healthy crown for years after local conspecifics have died(3, 54). Over the last 14 years, individuals meeting these criteria have been clonally propagated through grafting and subjected to greenhouse bioassays that provided evidence of the ability of some selected lingering ash trees to mount defensive responses against EAB (24). Although there is evidence of multiple types of defenses, this work is focused on EAB egg bioassays (described below) to assess host defenses that result in larval mortality. Clonal replicates of lingering green ash genotypes, some used as parents in this study, consistently kill more early instar larvae (35 to 50 percent) than the susceptible green ash controls (0 to 10 percent) (24).

### Plant Material

Plant material was comprised of 97 two-year-old potted *F. pennsylvanica* seedlings reared in an outdoor growing area, then transferred into an environmentally controlled greenhouse in the spring of the treatment year to allow acclimatization prior to the start of the EAB treatment. The individuals tested were generated by controlled cross-pollinations of lingering or susceptible green ash to produce full sibling families of known parentage. Individuals belonged to one of three families: Pe-C (21 individuals, susceptible parentage Pe-97 x “Summit”), Pe-Y (42 individuals, lingering parentage, Pe-53 x Pe-56), or Pe-Z (35 individuals, lingering parentage Pe-53 x Pe-59). Both susceptible parents, (Pe-97) and the cultivar “Summit”, had susceptible phenotypes in egg bioassays and did not persist on the landscape after the arrival of EAB. “Summit”, in particular, has been proven susceptible in our egg bioassay (16 replications), in common garden studies (55), and by its rapid demise under natural EAB infestation in city streets and parks (16).

### Emerald Ash Borer Resistance Bioassays

EAB eggs were raised and prepared as described in Koch et al 2015 (24). Twelve eggs were applied to each tree at a density of 400 eggs per square meter, as previously described (24). Eight weeks after eggs were applied, larval galleries were carefully dissected, starting at the entry hole, and followed to determine the outcome of each larva that successfully hatched and entered the tree. Larvae were designated as alive, tree-killed (killed by a host defense response), or dead by other means such as parasitism, cannibalism, or fungal infection. The proportion of tree-killed larvae was calculated based on the total number of larvae that hatched and entered the tree. One-way ANOVA and Tukey-Kramer multiple comparison tests were used to analyze the performance of families Pe-Y, Pe-Z and Pe-C.

### Metabolite analyses of *F. pennsylvanica* woody tissue by UHPLC-MS

Trees were destructively sampled to collect tissue for metabolite analyses eight weeks after eggs were placed, during phenotyping. The entire stem, 2.5 cm above the highest EAB larval galleries, was collected and stored immediately on dry ice, before being transferred to −80°C storage. This ensured the collection of the vascular cambium, the cork cambium, the phloem, and the ray parenchyma. All samples were ground under liquid nitrogen in a Spex Sample Prep freezer mill and stored at −80°C prior to extraction

For each sample, 1 g of frozen powdered plant tissues was extracted in 10 ml of acetonitrile/isopropanol/water (3:3:2) containing 1.00 mM telmisartan (internal standard) and 0.01% formic acid and incubated in the dark at 4°C for 24 h. samples were then centrifuged at 4°C and 10,000g for 10 minutes, supernatants were transferred to fresh tubes, and 50:1 diluted aliquots were prepared by adding deionized water. An additional aliquot of undiluted extracted sample has been archived at −80 °C.

UHPLC/MS analyses were performed using a Shimadzu LC-20AD ternary pump coupled to a SIL-5000 autosampler, column oven, and Waters Xevo G2-XS QTof mass spectrometer equipped with an electrospray ionization source. The operation parameters for the positive-ion mode analyses are as previously detailed(56). A 10-µL volume of each diluted extract was analyzed using a 20-minute gradient method on an Ascentis Express C18UHPLC column (2.1×100mm, 2.7µm) with mobile phases consisting of 10 mM ammonium formate in water, adjusted to pH 2.8 with formic acid (solvent A) and acetonitrile (solvent B). The 20-min method gradient was as follows: 1% B at 0.00 to 1.00 min, then step to 5% B at 1.01 min, linear gradient to 25% B at 8.00 min, then a linear gradient to 75% B at 12.50 min, another linear gradient to 98% B at 15.00 min, and a hold at 98% B until 18.00 min, a step to 1% B at 18.01 min, and a hold at 1% B until 20.00 min.

Analyte samples were injected in a randomized order while process blank and quality control samples were injected at regular intervals. All calculated peak areas were normalized to the peak area for the internal standard telmisartan utilizing Progenesis QI v2.4software (Nonlinear Dynamics Ltd., Newcastle, UK). Standards of oleuropein, apigenin and salidroside were purchased from Sigma Aldrich, prepared in the extraction solvent, and run at 5 µg/mL.

### Untargeted Metabolomics Data Processing

For untargeted metabolomic analysis, data were initially processed using Progenesis software. Leucine enkephalin lockmass correction (*m/z* 556.2766) was applied during run importation and all runs were aligned to retention times of a bulk pool run automatically selected by the software from a selection of QC samples. Peak picking and deconvolution was conducted as previously described (57). After deconvolution, 1,278 compound ions remained. To remove features from the dataset introduced by solvents, glassware, or instrumentation and to remove lipids, several filters were applied to the 1,278 compound ions remaining after deconvolution. Concentrations of each feature were normalized to the internal telmisartan standard (*m/z = 515*.*2448*). Compounds with the highest mean abundance in process blank samples, maximum abundance less than 0.1% of the most abundant compound in the dataset, or retention times greater than 16 minutes were excluded from the dataset. This reduced the total number of metabolic features to 323. Further analysis and statistical comparisons of compound signals extracted by Progenesis QI software was executed using EZinfo v3.0.2 software (Umetrics, Umeå, Sweden).

One way analysis of variance (ANOVA) tests were used to assess significance between each pairwise comparison for individual metabolic features in the seven following contrasts: Family C: UNI vs LLK ; Family Y: UNI vs LLK, UNI vs HLK, LLK vs HLK; Family Z: UNI vs LLK, UNI vs HLK, LLK vs HLK. Features that were significant (p < 0.05) were included in pairwise orthogonal partial least squares discriminant analysis (OPLS-DA) and principal component analysis (PCA) analyses (Table S1). OPLS-DAs and PCAs were run using pareto scaling. To prevent overfitting of our model we performed a threefold cross validation on our data and took the averages as the performance of our model. For all metabolic features extracted with Progenesis QI and used in downstream analyses with EZinfo, spectra were processed using MassLynx v4.2 software (Waters Corporation, Milford, MA, USA) as previously detailed (57) (Table S2).

Of the metabolites considered, thirty-two had significance in more than one family, or had a previously proposed purpose and were annotated. Annotation of the electrospray ionization tandem mass spectrometry (ESI-MS/MS) data relied on comparisons with MS/MS databases such as the Massbank of North America as well as previous studies and purchased standards. The confidence levels in the metabolite annotation were following recommendations of the Metabolomics Standards Initiative (48). The quantities present in individual tissue extracts were too small for complete structure elucidation.

## Acknowledgements

The authors thank Warren Chatwin and Christina Murray for their helpful comments on the manuscript. The authors thank Aletta Doran, Julia Wolf, Gavin Nupp, Miranda McKibben, and Jarod Sanchez for their work propagating and maintaining the study trees and their assistance conducting the EAB resistance bioassays. The authors also thank Patrick Cunniff, Brandon Chou, Kingsley Owusu Otoo and Julie Huston for assistance in collecting and organizing tissue samples and managing logistics. J.R-S acknowledges support from USDA-USFS APHIS grants 18-IA-11242316-105 and 20-JV-11242303-050. J.R-S also acknowledges support from the Tree Fund Foundation, Tree Fund grant 18-JD-01. R.K.S. acknowledges support from NIH training grant T32GM075762. JK acknowledges support from USDA APHIS 18-IA-11242316-105, Michigan Invasive Species Grant Program grant IS18-119, the Commonwealth of Pennsylvania Department of Conservation and Natural Resources Bureau of Forestry 18-CO-11242316-014, and the U.S Forest Service Special Technology Development Program grant NA-2017-01. A.D.J. acknowledges support from Michigan AgBioResearch through the USDA National Institute of Food and Agriculture, Hatch project number MICL02474, and USDA-USFS grant 20-JV-11242303-050.

## Data Availability

The data that support the findings of this study are available upon request from the corresponding author. The raw data will also be submitted to MetaboLights or similar repository.

## References

1. K. M. Potter, M. E. Escanferla, R. M. Jetton, G. Man, Important Insect and Disease Threats to United States Tree Species and Geographic Patterns of Their Potential Impacts. Forests 10, 304 (2019).

2. D. G. McCullough, Challenges, tactics and integrated management of emerald ash borer in North America. Forestry: An International Journal of Forest Research 10.1093/forestry/cpz049, cpz049 (2019).

3. K. S. Knight et al. (2012) Dynamics of surviving ash (Fraxinus spp.) populations in areas long infested by emerald ash borer (Agrilus planipennis). in Proceedings of the fourth international workshop on the genetics of host-parasite interactions in forestry: Disease and insect resistance in forest trees (Pacific Southwest Research Station, Forest Service, U.S. Department of Agriculture, Albany, CA), pp 143–152.

4. J. L. Koch, D. W. Carey, M. E. Mason, T. M. Poland, K. S. Knight, Intraspecific variation in Fraxinus pennsylvanica responses to emerald ash borer (Agrilus planipennis). New Forests 46, 995–1011 (2015).

5. C. F. Miniat et al., “Impacts of Invasive Species on Forest and Grassland Ecosystem Processes in the United States” in Invasive Species in Forests and Rangelands of the United States, T. M. Poland et al., Eds. (Springer International Publishing, Cham, 2021), pp. 41–55.

6. W. R. L. Anderegg et al., Climate-driven risks to the climate mitigation potential of forests. Science 368, eaaz7005 (2020).

7. J. A. Hicke et al., Effects of biotic disturbances on forest carbon cycling in the United States and Canada. Glob Change Biol 18, 7–34 (2012).

8. K. Hoover, A. A. Riddle, Forest carbon primer. Congressional Research Service: Washington, DC, USA (2020).

9. K. F. Kovacs et al., Cost of potential emerald ash borer damage in U.S. communities, 2009–2019. Ecological Economics 69, 569–578 (2010).

10. J. E. Aukema et al., Economic impacts of non-native forest insects in the continental United States. PLoS one 6, e24587 (2011).

11. C. J. A. Bradshaw et al., Massive yet grossly underestimated global costs of invasive insects. Nature Communications 7, 12986 (2016).

12. G. M. Lovett et al., Nonnative forest insects and pathogens in the United States: Impacts and policy options. Ecol Appl 26, 1437–1455 (2016).

13. F. Krist Jr et al., National Insect and Disease Forest Risk Assessment: 2013-2027. US Department of Agriculture. FHTET-14-01 (2014).

14. K. Potter, M. Escanferla, R. Jetton, G. Man, Important Insect and Disease Threats to United States Tree Species and Geographic Patterns of Their Potential Impacts. Forests 10, 304 (2019).

15. K. M. Potter, M. E. Escanferla, R. M. Jetton, G. Man, B. S. Crane, Prioritizing the conservation needs of United States tree species: Evaluating vulnerability to forest insect and disease threats. Global Ecology and Conservation 18, e00622 (2019).

16. T. M. Poland, D. G. McCullough, Emerald ash borer: invasion of the urban forest and the threat to North America’s ash resource. Journal of Forestry 104, 118–124 (2006).

17. D. A. Herms, D. G. McCullough, Emerald ash borer invasion of North America: history, biology, ecology, impacts, and management. Annual review of entomology 59, 13–30 (2014).

18. G. Popkin, Rising from the ashes Science 370, 756–759 (2020).

19. B. B. Hanberry, Rise of Fraxinus in the United States between 1968 and 20131. The Journal of the Torrey Botanical Society 141, 242–249 (2014).

20. K. S. Knight, J. P. Brown, R. P. Long, Factors affecting the survival of ash (Fraxinus spp.) trees infested by emerald ash borer (Agrilus planipennis). Biological Invasions 15, 371–383 (2013).

21. Anonymous (IUCN 2021. The IUCN Red List of Threatened Species. Version 2021-1. https://www.iucnredlist.org. Downloaded on 6/27/2021.

22. R. A. Sniezko et al., Proceedings of the fourth international workshop on the genetics of host-parasite interactions in forestry: Disease and insect resistance in forest trees. Gen. Tech. Rep. PSW-GTR-240. Albany, CA: Pacific Southwest Research Station, Forest Service, US Department of Agriculture. 372 p 240 (2012).

23. J. L. Koch et al. (2012) Breeding strategies for the development of emerald ash borer-resistant North American ash. in In: Sniezko, Richard A.; Yanchuk, Alvin D.; Kliejunas, John T.; Palmieri, Katharine M.; Alexander, Janice M.; Frankel, Susan J., tech. coords. Proceedings of the fourth international workshop on the genetics of host-parasite interactions in forestry: Disease and insect resistance in forest trees. Gen. Tech. Rep. PSW-GTR-240. Albany, CA: Pacific Southwest Research Station, Forest Service, US Department of Agriculture. pp. 235–239., pp 235-239.

24. J. Koch, D. Carey, M. Mason, T. Poland, K. Knight, Intraspecific variation in Fraxinus pennsylvanica responses to emerald ash borer (Agrilus planipennis). New Forests 46, 995–1011 (2015).

25. J. Romero-Severson, J. L. Koch (2017) Saving green ash. in Proceedings of Workshop on Gene Conservation of Tree Species-Banking on the Future, May 2016, pp 102–110.

26. R. A. Sniezko, J. Koch, Breeding trees resistant to insects and diseases: putting theory into application. Biological Invasions 19, 3377–3400 (2017).

27. M.-L. Desprez-Loustau et al., An evolutionary ecology perspective to address forest pathology challenges of today and tomorrow. Annals of Forest Science 73, 45–67 (2016).

28. K. J. K. Gandhi, D. A. Herms, Direct and indirect effects of alien insect herbivores on ecological processes and interactions in forests of eastern North America. Biological Invasions 12, 389–405 (2010).

29. A. M. Mech et al., Evolutionary history predicts high-impact invasions by herbivorous insects. Ecology and evolution 9, 12216–12230 (2019).

30. C. C. Pike, J. Koch, C. D. Nelson, Breeding for Resistance to Tree Pests: Successes, Challenges, and a Guide to the Future. Journal of Forestry 119, 96–105 (2020).

31. R. A. Sniezko, J.-J. Liu, Prospects for developing durable resistance in populations of forest trees. New Forests, 1–17 (2021).

32. J. L. Koch, R. L. Heyd, Battling beech bark disease: establishment of beech seed orchards in Michigan. Newsletter of the Michigan Entomological Society. 58 (1&2): 11–14. 58, 11-14 (2013).

33. C. C. Pike et al., Improving the resistance of eastern white pine to white pine blister rust disease. Forest ecology and management 423, 114–119 (2018).

34. J. G. Whitehill et al., Interspecific comparison of constitutive ash phloem phenolic chemistry reveals compounds unique to Manchurian ash, a species resistant to emerald ash borer. Journal of Chemical Ecology 38, 499–511 (2012).

35. J. G. A. Whitehill, C. Rigsby, D. Cipollini, D. A. Herms, P. Bonello, Decreased emergence of emerald ash borer from ash treated with methyl jasmonate is associated with induction of general defense traits and the toxic phenolic compound verbascoside. Oecologia 176, 1047–1059 (2014).

36. C. Villari, D. A. Herms, J. G. Whitehill, D. Cipollini, P. Bonello, Progress and gaps in understanding mechanisms of ash tree resistance to emerald ash borer, a model for wood-boring insects that kill angiosperms. New Phytologist 209, 63–79 (2016).

37. L. J. Kelly et al., Convergent molecular evolution among ash species resistant to the emerald ash borer. Nature ecology & evolution 4, 1116–1128 (2020).

38. R. Enderle, J. Stenlid, R. Vasaitis, An overview of ash (Fraxinus spp.) and the ash dieback disease in Europe. CAB Rev 14, 1–12 (2019).

39. E. S. A. Sollars et al., Genome sequence and genetic diversity of European ash trees. Nature 541, 212–216 (2017).

40. M. Nemesio-Gorriz et al., Canditate metabolites for ash dieback tolerance in Fraxinus excelsior. Journal of Experimental Botany 71, 6074–6083 (2020).

41. L. McKinney et al., The ash dieback crisis: genetic variation in resistance can prove a long-term solution. Plant Pathology 63, 485–499 (2014).

42. X. López-Goldar et al., Inducibility of plant secondary metabolites in the stem predicts genetic variation in resistance against a key insect herbivore in maritime pine. Frontiers in Plant Science 9, 1651 (2018).

43. X. López-Goldar et al., Genetic variation in the constitutive defensive metabolome and its inducibility are geographically structured and largely determined by demographic processes in maritime pine. Journal of Ecology 107, 2464–2477 (2019).

44. M. Volf, J. Hrcek, R. Julkunen-Tiitto, V. Novotny, To each its own: differential response of specialist and generalist herbivores to plant defence in willows. Journal of Animal Ecology 84, 1123–1132 (2015).

45. K. F. Raffa, E. B. Smalley, Interaction of pre-attack and induced monoterpene concentrations in host conifer defense against bark beetle-fungal complexes. Oecologia 102, 285–295 (1995).

46. B. Worley, R. Powers, PCA as a practical indicator of OPLS-DA model reliability. Current Metabolomics 4, 97–103 (2016).

47. S. Chakraborty et al., Effects of water availability on emerald ash borer larval performance and phloem phenolics of Manchurian and black ash. Plant, Cell & Environment 37, 1009–1021 (2014).

48. L. W. Sumner et al., Proposed minimum reporting standards for chemical analysis. Metabolomics 3, 211–221 (2007).

49. J. D. Sidda et al., Diversity of secoiridoid glycosides in leaves of UK and Danish ash provide new insight for ash dieback management. Scientific reports 10, 1–12 (2020).

50. D. Falconer, T. Mackay, Introduction to quantitative genetics 4th edition. Harlow, UK: Longmans (1996).

51. M. Huff et al., A high-quality reference genome for Fraxinus pennsylvanica for ash species restoration and research. Molecular ecology resources (2021).

52. J. G. Whitehill, C. Rigsby, D. Cipollini, D. A. Herms, P. Bonello, Decreased emergence of emerald ash borer from ash treated with methyl jasmonate is associated with induction of general defense traits and the toxic phenolic compound verbascoside. Oecologia 176, 1047–1059 (2014).

53. M. C. Schuman, I. T. Baldwin, The layers of plant responses to insect herbivores. Annual review of entomology 61, 373–394 (2016).

54. K. C. Steiner, L. E. Graboski, K. S. Knight, J. L. Koch, M. E. Mason, Genetic, spatial, and temporal aspects of decline and mortality in a Fraxinus provenance test following invasion by the emerald ash borer. Biological Invasions 21, 3439–3450 (2019).

55. E. J. Rebek, D. A. Herms, D. R. Smitley, Interspecific Variation in Resistance to Emerald Ash Borer (Coleoptera: Buprestidae) Among North American and Asian Ash (Fraxinus spp.). Environmental Entomology 37, 242–246 (2008).

56. D. B. Lybrand, T. M. Anthony, A. D. Jones, R. L. Last, An integrated analytical approach reveals trichome acylsugar metabolite diversity in the wild tomato Solanum pennellii. Metabolites 10, 401 (2020).

57. R. Sadre et al., Metabolite diversity in alkaloid biosynthesis: a multilane (diastereomer) highway for camptothecin synthesis in Camptotheca acuminata. The Plant Cell 28, 1926–1944 (2016).

