## Supplemental Figures 1-3, Supplemental Tables 1-2 for "Profiles of secoiridoids and alkaloids in tissue of susceptible and resistant green ash progeny reveal patterns of induced responses to emerald ash borer in *Fraxinus pennsylvanica*"

**This PDF file includes:**

Figures S1-S3

Tables S1-S2

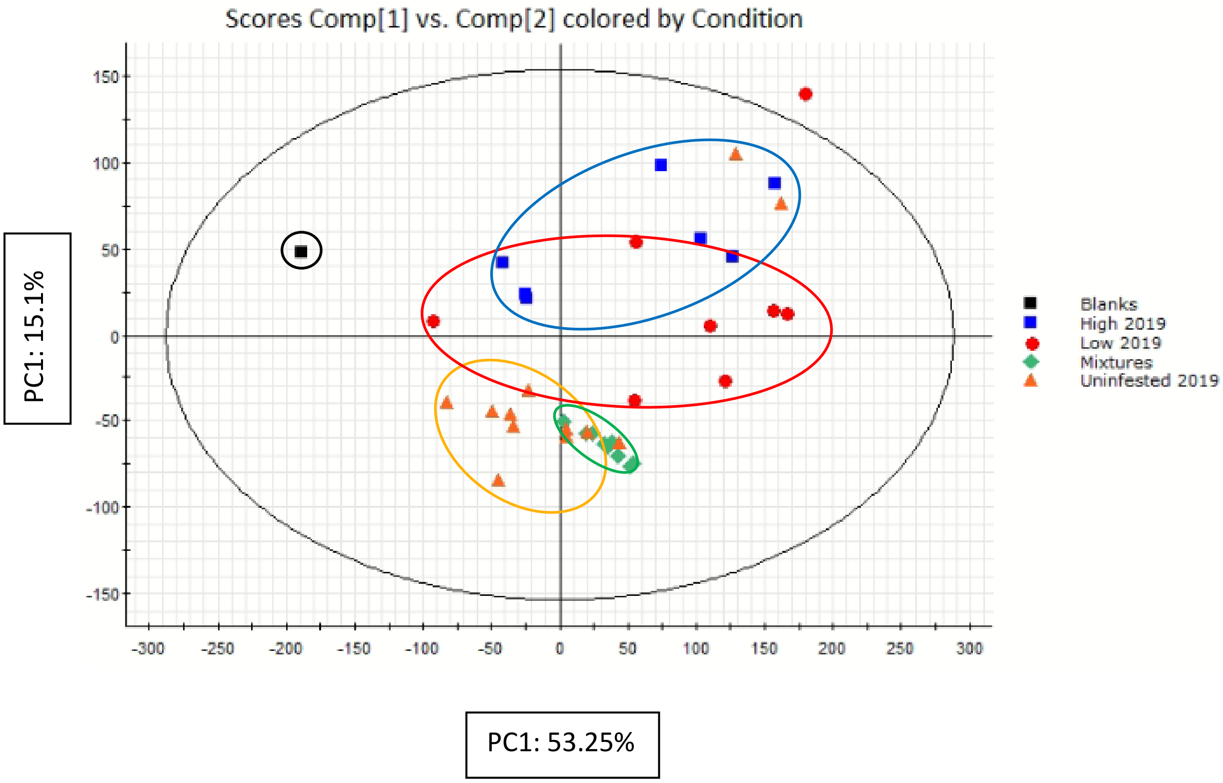


Figure S1: PCA of Blanks, controls (mixtures), low larval kill, high larval kill and uninfested individuals from family Pe-Y. 194 features were utilized. Distinct groups were visualized.

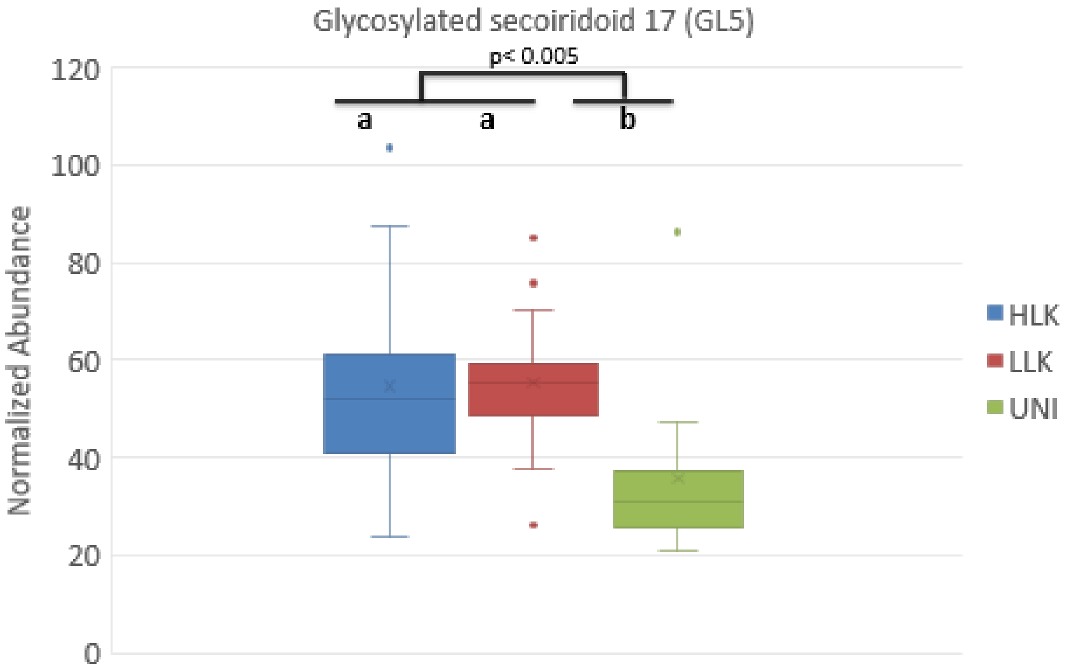


Figure S2: Relative concentrations of GL5 in High Larval Kill (HLK) Low Larval Kill (LLK) and Uninfested (UNI) groups in family Pe-Y.

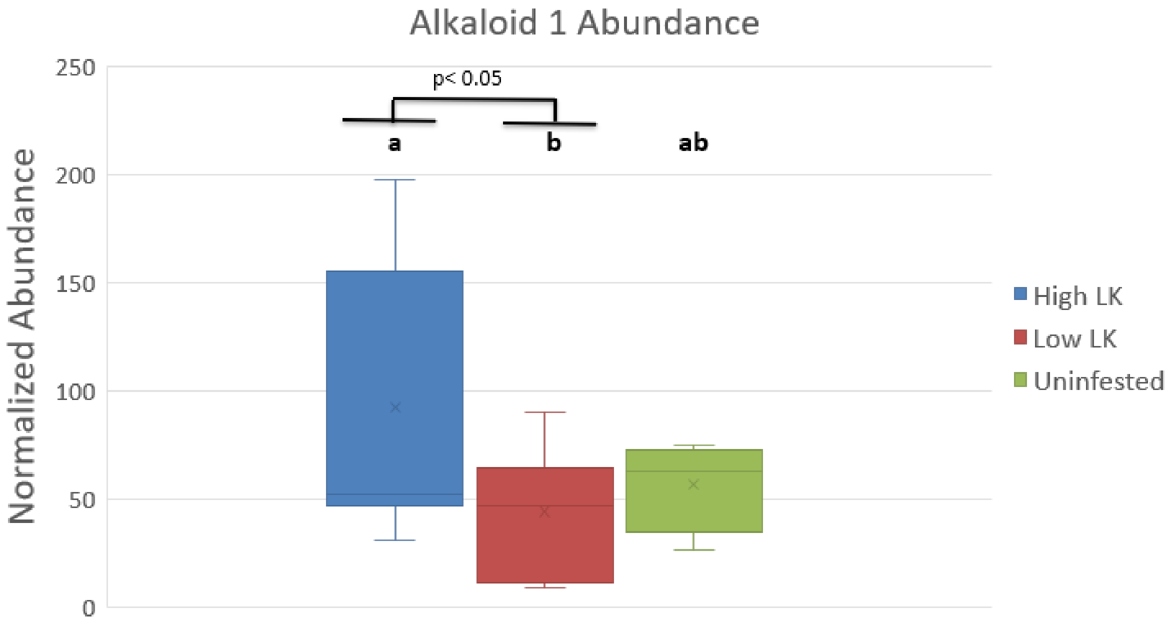


Figure S3: Relative concentrations of Alkaloid 1 in High Larval Kill (HLK) Low Larval Kill (LLK) and Uninfested (UNI) groups in family Pe-Z.

Table S1: Lists of features included in each pairwise comparison, and which group their concentrations were higher in. (letter indicates group that feature is highest in (p<.05))

| Retention time (min) | m/z | (Y) 2019 LvU .05 Main Peaks |
|---|---|---|
| 0.83 | 200.11 | L |
| 0.93 | 325.11 | U |
| 0.93 | 684.25 | U |
| 0.95 | 505.17 | U |
| 0.97 | 136.06 | L |
| 1.26 | 240.12 | L |
| 5.16 | 365.12 | L |
| 5.57 | 584.22 | L |
| 5.73 | 234.08 | L |
| 5.95 | 151.04 | L |
| 5.98 | 422.17 | L |
| 6.20 | 561.19 | L |
| 6.65 | 282.17 | L |
| 6.71 | 427.12 | L |
| 6.73 | 700.28 | L |
| 7.28 | 545.20 | L |
| 7.43 | 554.22 | L |
| 7.62 | 543.18 | L |
| 7.89 | 642.24 | L |
| 8.26 | 970.34 | L |
| 8.39 | 1063.38 | L |
| 8.40 | 705.28 | L |
| 8.43 | 704.28 | L |
| 8.55 | 163.04 | L |
| 8.61 | 836.32 | L |
| 8.65 | 596.24 | U |
| 8.70 | 235.10 | L |
| 8.74 | 361.13 | L |
| 8.78 | 970.34 | L |
| 8.81 | 731.29 | L |
| 8.83 | 704.28 | L |
| 8.87 | 507.19 | L |
| 8.94 | 601.19 | U |
| 8.96 | 519.19 | U |
| 9.20 | 325.09 | L |
| 9.35 | 700.28 | L |

| Retention time (min) | m/z | (C) 2019 .05 LvU Major Peaks |
|---|---|---|
| 0.83 | 200.11 | L |
| 0.95 | 505.17 | U |
| 1.28 | 325.14 | L |
| 1.35 | 503.12 | L |
| 4.52 | 487.14 | L |
| 4.55 | 431.15 | L |
| 4.96 | 167.07 | L |
| 5.16 | 365.12 | L |
| 5.44 | 439.16 | L |
| 5.83 | 425.14 | L |
| 5.92 | 341.14 | L |
| 6.12 | 196.10 | L |
| 6.20 | 561.19 | L |
| 6.77 | 439.16 | L |
| 6.95 | 591.20 | L |
| 7.28 | 545.20 | L |
| 7.43 | 554.22 | L |
| 7.54 | 137.06 | L |
| 7.62 | 543.18 | L |
| 7.63 | 591.20 | L |
| 7.80 | 573.19 | L |
| 8.33 | 543.18 | L |
| 8.40 | 165.06 | L |
| 8.43 | 704.28 | L |
| 8.61 | 836.32 | L |
| 8.67 | 598.25 | L |
| 8.68 | 568.24 | L |
| 8.74 | 558.22 | L |
| 8.75 | 586.25 | L |
| 8.78 | 970.34 | L |
| 8.83 | 704.28 | L |
| 9.35 | 700.28 | L |
| 9.51 | 769.27 | L |
| 9.88 | 764.32 | L |
| 9.95 | 542.23 | L |
| 9.95 | 569.23 | L |

| Retention time (min) | m/z | (Z) 2019 LvU .05 Main Peaks |
|---|---|---|
| 0.83 | 175.12 | U |
| 0.85 | 104.11 | U |
| 0.95 | 505.17 | U |
| 1.26 | 452.18 | U |
| 5.57 | 584.22 | L |
| 5.98 | 422.17 | L |
| 6.05 | 165.05 | L |
| 6.73 | 700.28 | L |
| 6.82 | 165.05 | L |
| 6.93 | 804.29 | L |
| 7.28 | 760.30 | U |
| 7.89 | 642.24 | L |
| 8.04 | 640.22 | L |
| 8.08 | 361.16 | U |
| 8.67 | 598.25 | U |
| 8.72 | 357.13 | U |
| 9.75 | 342.13 | L |
| 9.88 | 764.32 | U |
| 9.95 | 569.23 | L |
| 10.27 | 542.23 | L |
| 10.85 | 801.27 | U |
| 10.95 | 404.17 | U |
| 11.08 | 833.30 | U |
| 11.13 | 581.20 | U |
| 11.23 | 446.18 | U |

| | | |
|---|---|---|
| 9.75 | 388.17 | L |
| 9.88 | 840.29 | L |
| 9.90 | 446.22 | L |
| 9.95 | 542.23 | L |
| 9.95 | 569.23 | L |
| 10.10 | 769.26 | L |
| 10.23 | 1090.39 | L |
| 10.27 | 542.23 | L |
| 10.38 | 1164.41 | L |
| 10.40 | 577.19 | L |
| 10.78 | 928.34 | L |
| 11.02 | 734.26 | L |
| 11.32 | 726.33 | L |

| | | |
|---|---|---|
| 9.97 | 331.12 | L |
| 10.29 | 345.13 | L |
| 10.38 | 607.21 | L |
| 10.42 | 944.34 | L |
| 10.78 | 928.34 | L |
| 11.78 | 238.14 | L |
| 14.30 | 411.09 | L |

| Retention time (min) | m/z | (Y)2019 HvU .05 major peaks |
|---|---|---|
| 0.83 | 175.12 | U |
| 1.26 | 282.13 | H |
| 1.28 | 424.18 | U |
| 4.52 | 487.14 | U |
| 4.61 | 282.13 | H |
| 5.16 | 365.12 | H |
| 5.57 | 584.22 | H |
| 5.98 | 165.05 | H |
| 6.62 | 663.19 | H |
| 6.64 | 325.09 | H |
| 6.65 | 282.17 | H |
| 7.28 | 265.11 | U |
| 7.28 | 760.30 | U |
| 7.30 | 205.09 | U |
| 7.40 | 324.18 | H |
| 7.43 | 554.22 | H |
| 7.63 | 591.20 | U |
| 7.67 | 219.10 | U |

| Retention time (min) | m/z | (Z) 2019 HvU .05 major peaks |
|---|---|---|
| 0.85 | 104.11 | U |
| 0.93 | 846.30 | H |
| 1.26 | 452.18 | U |
| 1.35 | 503.12 | U |
| 5.57 | 584.22 | H |
| 5.98 | 422.17 | H |
| 6.05 | 165.05 | H |
| 6.95 | 802.27 | H |
| 7.28 | 760.30 | U |
| 7.54 | 137.06 | U |
| 8.00 | 519.25 | U |
| 8.74 | 558.22 | U |
| 8.85 | 265.11 | U |
| 8.94 | 601.19 | U |
| 9.33 | 546.25 | U |
| 9.88 | 764.32 | U |
| 9.92 | 401.12 | U |
| 9.95 | 569.23 | H |

| Retention time (min) | m/z | (Y) 2019 HvL .05 Main Peaks |
|---|---|---|
| 0.85 | 104.11 | L |
| 1.26 | 282.13 | H |
| 1.28 | 424.18 | L |
| 4.32 | 457.13 | L |
| 4.61 | 282.13 | H |
| 4.89 | 323.11 | H |
| 5.50 | 395.13 | L |
| 5.50 | 767.27 | L |
| 6.84 | 561.19 | L |
| 7.23 | 547.21 | L |
| 7.25 | 570.29 | L |
| 7.28 | 265.11 | L |
| 7.28 | 760.30 | L |
| 7.30 | 205.09 | L |
| 7.63 | 591.20 | L |
| 7.67 | 449.11 | L |
| 7.69 | 545.20 | L |
| 7.82 | 589.18 | L |

| | | |
|---|---|---|
| 7.95 | 575.21 | U |
| 7.97 | 591.19 | U |
| 8.00 | 519.25 | U |
| 8.10 | 467.15 | U |
| 8.32 | 647.19 | U |
| 8.65 | 596.24 | U |
| 8.68 | 589.17 | U |
| 8.70 | 603.20 | U |
| 8.72 | 371.15 | U |
| 8.74 | 531.27 | U |
| 8.81 | 731.29 | H |
| 8.83 | 704.28 | H |
| 9.20 | 325.09 | H |
| 9.33 | 546.25 | U |
| 9.35 | 700.28 | H |
| 9.88 | 764.32 | U |
| 9.95 | 569.23 | H |
| 9.95 | 542.23 | H |
| 9.97 | 870.30 | U |
| 9.99 | 501.23 | U |
| 10.38 | 1164.41 | H |
| 10.75 | 271.06 | U |
| 10.97 | 151.04 | H |
| 12.50 | 316.28 | U |
| 12.79 | 318.30 | U |
| 15.20 | 600.42 | U |

| | | |
|---|---|---|
| 9.99 | 501.23 | U |
| 10.27 | 542.23 | H |
| 11.23 | 446.18 | U |

| | | |
|---|---|---|
| 7.89 | 1266.44 | L |
| 7.95 | 575.21 | L |
| 7.97 | 591.19 | L |
| 8.10 | 467.15 | L |
| 8.26 | 970.34 | L |
| 8.32 | 647.19 | L |
| 8.40 | 165.06 | L |
| 8.43 | 704.28 | L |
| 8.61 | 836.32 | L |
| 8.70 | 603.20 | L |
| 8.72 | 371.15 | L |
| 8.74 | 558.22 | L |
| 8.75 | 586.25 | L |
| 8.78 | 970.34 | L |
| 8.81 | 387.14 | L |
| 9.55 | 776.29 | L |
| 9.83 | 581.24 | L |
| 9.88 | 764.32 | L |
| 9.94 | 1028.36 | L |
| 9.97 | 870.30 | L |
| 10.23 | 1090.39 | L |
| 10.40 | 359.15 | L |
| 10.75 | 271.06 | L |
| 10.75 | 713.20 | L |
| 11.02 | 734.26 | L |

| Retention time (min) | m/z | (Z) 2019 HvL .05 Main Peaks |
|---|---|---|
| 1.26 | 240.12 | H |
| 4.61 | 282.13 | H |
| 6.819483 | 165.05 | L |
| 7.668317 | 219.10 | L |
| 7.798967 | 189.09 | L |
| 8.2374 | 533.16 | L |
| 8.74 | 558.22 | L |
| 8.90225 | 572.23 | L |
| 11.32 | 726.33 | L |

Table S2: Performance of OPLS-DA models

| | number of features used | % individuals correctly assigned | % individuals incorrectly assigned | % individuals not assigned by model |
|---|---|---|---|---|
| (Pe-C) LLK v UNI | 34 | 79 | 5 | 16 |
| (Pe-Y) LLK v UNI | 49 | 83 | 11 | 6 |
| (Pe-Z) LLK v UNI | 25 | 83 | 8 | 8 |
| (Pe-Y) HLK v UNI | 44 | 73 | 7 | 20 |
| (Pe-Z) HLK v UNI | 35 | 87 | 0 | 13 |
| (Pe-Y) HLK v LLK | 43 | 71 | 14 | 14 |
| (Pe-Z) HLK v LLK | 9 | 76 | 6 | 18 |
